# *In vitro* membrane remodelling by ESCRT-II/ESCRT-III is regulated by negative feedback from membrane tension

**DOI:** 10.1101/438481

**Authors:** Andrew Booth, Christopher J. Marklew, Barbara Ciani, Paul A. Beales

## Abstract

Artificial cells can shed new light on the molecular basis for life and hold potential for new chemical technologies. Inspired by how nature dynamically regulates its membrane compartments, we aim to repurpose the endosomal sorting complex required for transport (ESCRT) to generate complex membrane architectures as suitable scaffolds for artificial cells. Purified ESCRT-III components perform topological transformations on giant unilamellar vesicles (GUVs) to create complex “vesicles-within-a-vesicle” architectures resembling the compartmentalisation in eukaryotic cells. Thus far, the proposed mechanisms for this activity are based on how assembly and disassembly of ESCRT-III on the membrane drives deformation. Here we demonstrate the existence of a negative feedback mechanism from membrane mechanics that regulates ESCRT-III activity. ILV formation removes excess membrane area, increasing tension, which in turn suppresses downstream ILV formation. This mechanism for *in vitro* regulation of ESCRT-III activity may also have important implications for its *in vivo* functions.

## Introduction

The biological cell is fundamentally a highly complex chemical reactor, where compartmentalisation of its chemical processes is essential for complex functions.^1^ Eukaryotic organisms contain membrane-bound cellular sub-compartments, which allow distinct chemical environments to exist within the cell, such that otherwise incompatible chemistries can be maintained and utilised, e.g. ATP synthesis in the mitochondria and protein degradation in lysosomes.^2^ The manifold capabilities of a cell in the synthesis of complex biomolecules, sensing and response to its environment are all desirable functionalities to emulate within a synthetic system.^3^ Therefore chemical technologies that provide control over the formation of multicompartment membrane architectures are fundamental to artificial cell engineering.^4,5^

**Figure 1.**
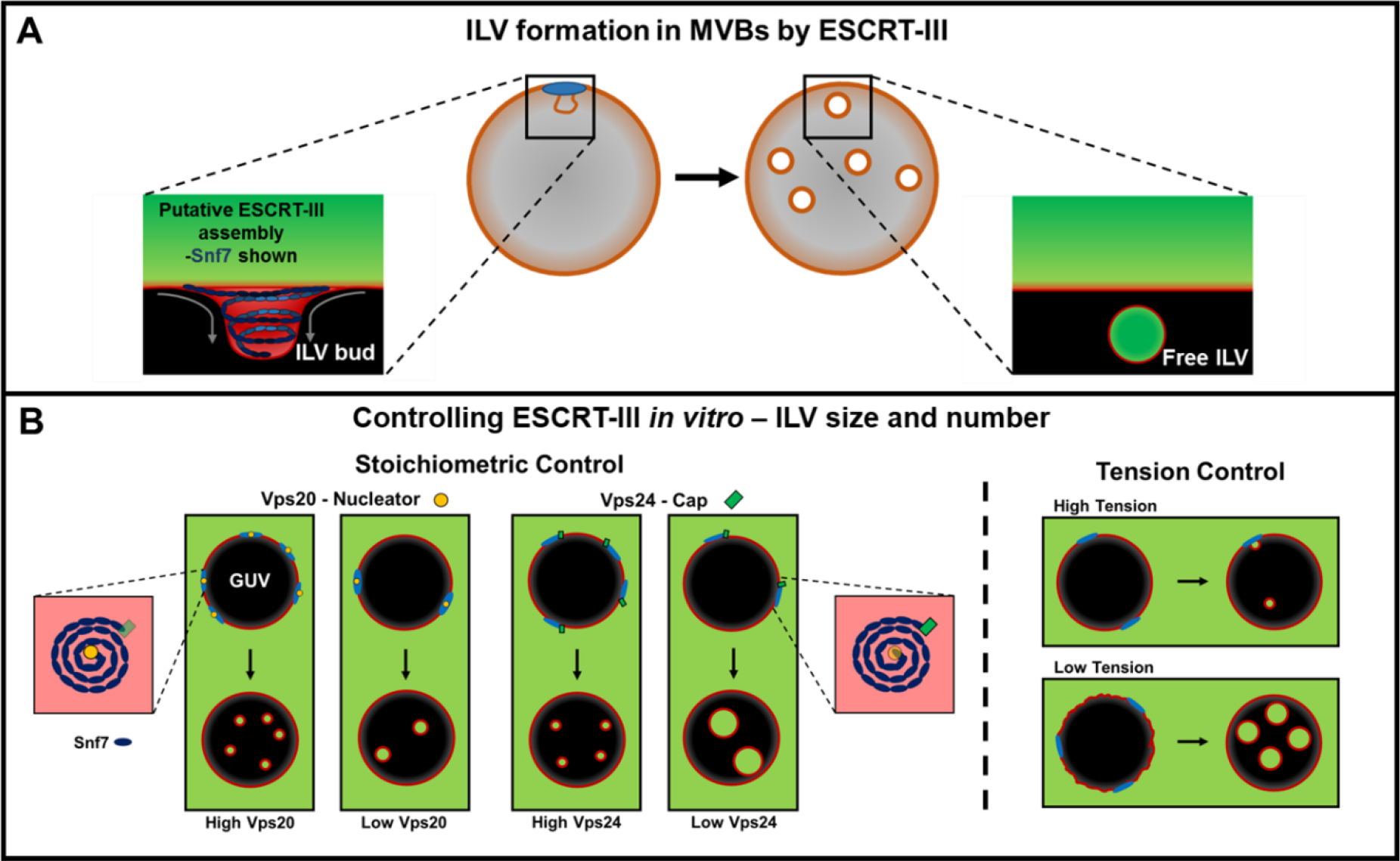
Experimental design rationale. **A**: ESCRT-III drives a number of cellular processes that generate new membrane-bound compartments. Notably, multi-vesicular body (MVB) genesis, this process that can be reconstituted *in vitro* using synthetic vesicles^6^ **B**: ESCRT-III may deform membranes by assembling into ordered supramolecular ‘springs’ (depicted as blue spirals). Protein stoichiometry and membrane tension were identified as variables that may allow finer control over the size and number of ILVs produced *in vitro.* Nucleation of Snf7 spiral filaments by Vps20 may provide a means to control the number of ILVs, while Vps24 may act as a terminator or ‘cap’ of Snf7 filament growth, limiting ILV size. High membrane tension may impede ILV formation by resisting membrane deformation, potentially impacting both the size and number of ILVs formed.

While microfluidic technologies have been developed that allow fixed multicompartment membrane architecture to be formed,^7,8^ natural cells can dynamically change their compartmentalised architectures. To emulate this dynamic nature within an artificial cell system, we take inspiration from biology by repurposing protein complexes *in vitro* that natively remodel membrane structures.^9^ The “vesicles-within-a-vesicle” architecture we aspire to recreate in an artificial cell bears resemblance to a cellular membrane-bound organelle, the multivesicular body (MVB). In eukaryotic cells, MVBs are formed by the encapsulation of biomolecular cargo into intraluminal vesicles (ILVs) within endosomes.^10^ This membrane remodelling event is performed by the ESCRT (endosomal complex required for transport) family of proteins.^11^ This activity lends itself to the possibility of using ESCRTs in the construction of multicompartment artificial cells *in vitro*.^12^

The human ESCRT complex drives multiple key membrane-remodelling processes, such as multivesicular body formation, cell division, exosome formation, HIV budding, plasma and nuclear envelope membrane repair.^13,14^ In yeast, ESCRT is responsible for cargo sorting in the endosomal pathway and also for nuclear pore complex quality control.^15^ The ESCRT complex responsible for transmembrane receptor sorting via the endosomal pathway comprises of ESCRT-0, ESCRT-I and ESCRT-II, which capture ubiquitinated transmembrane cargo at the late endosome membrane.^16^ The central membrane remodelling complex is ESCRT-III a highly conserved set of proteins in eukaryotes, and the AAA+ ATPase Vps4 (**Figure 1A**).^17,18^ In *Saccharomyces cerevisiae*, ESCRT-III consists of a core machinery comprised of the Vps20, Snf7, Vps24 and Vps2 subunits. Membrane budding and scission of vesicles mediated by ESCRT-III occurs with a unique topology, whereby the membrane is pushed away from the cytosol.^19^ Initial membrane deformation is performed by ESCRT-II, which recruits ESCRT-III subunits and drives the formation of ESCRT-III filaments on the membrane.^20^ *In vitro* studies on flat surface-supported membranes have shown that ESCRT-III forms spirals that are proposed to behave like springs, storing elastic energy, which deform the membrane upon release.^21^ The observation of this spiral structure informs the mechanical rationale for membrane deformation and scission in many of the proposed mechanisms where either a spiral,^22^ cone,^23^ or dome^24,25^ structure made by ESCRT-III stabilises and subsequently drives scission of the ILV bud neck. This process is made possible by the action of the ATPase Vps4, which has been shown to catalyse ESCRT-III activity by maintaining the dynamics of complex assembly and disassembly.^26^

ESCRT-II recruits the ESCRT-III subunit Vps20 to form a complex with affinity for regions of membrane curvature^27^. In turn, Vps20 nucleates the formation of Snf7 spirals that are capped at the growing end by a Vps24-Vps2 complex.^28^ Vps2 recruits the ATPase Vps4 through the interaction between its MIM (MIT-Interacting Motif) motif with the MIT (Microtubule Interacting and Transport) domain of Vps4.^29^ Intraluminal vesicle (ILV) formation by ESCRT-II/ESCRT-III can be reconstituted *in vitro* with a GUV-based bulk-phase uptake assay. The activity of a minimal ESCRT-III / Vps4 system in artificial vesicle systems was first demonstrated by Wollert *et al.*^6^ where ILVs were generated in GUV parent vesicles and observed by microscopy.

The reconstitution of ESCRT activity *in vitro* opens the possibility of achieving finer control of ILV formation in artificial systems. Therefore we initially set out to investigate the relative stoichiometry of ESCRT-III components to underpin their influence on the size and number of ILVs that form within individual GUVs (**Figure 1B**). Interestingly, our initial experiments reveal wide variation in ILV size and number between different GUVs within an individual experiment (**Figure 2A)**. Membrane mechanics are considered to be the likely source of these differences due to variability in tension across a single GUV population. Modulating membrane tension could therefore provide a further means to exert control over ILV formation and will also be investigated in this study.

Here we report on how changing the relative stoichiometry of the ESCRT-III subunits and the mechanical properties of the membrane regulate the size and number of ILVs formed in a GUV-based bulk-phase uptake assay (**Figure 1B**). These findings reveal a novel mechanism that regulates ESCRT activity *in vitro* and provide further insight into how they may function *in vivo*.

## Results

### ‘Passive’ and ‘active’ membrane budding and scission by ESCRT

We have quantified ESCRT activity using the bulk phase uptake assay as described by Wollert and Hurley,^6^ introducing a systematic variation in the ratio of key ESCRT-III in the reaction. In this assay, GUVs with an ‘endosome-like’ lipid composition^6^ are incubated with ESCRT-II, and the core ESCRT-III subunits Vps20, Vps24, Vps2 and Snf7. Prior to the addition of proteins, a membrane-impermeable, fluorescent dextran (M_r_ ~ 5 kDa, Cascade blue labelled) is added to the GUV suspension to establish whether newly formed ILVs have encapsulated bulk phase (**Figure 2A**). Confocal microscopy (**Figure 2B**) is used to assess the number of ILVs and the observed diameter of each ILV is recorded (observed diameter may not be the true maximum diameter of the ILV). Only ILVs visibly containing dextran fluorescence are counted, as this confirms their formation following addition of the proteins. The number of ILVs and the volume of the GUV lumens in the image is then used to calculate the number of ILVs per unit volume, the unit volume being taken as that of an average 20 μm diameter GUV, hence ‘ILVs per GUV volume equivalent’. Controls containing only osmotically balanced buffer and fluorescent dextran give a minimal background level of intraluminal vesicles formed within the experimental period (see ‘no protein control’ in figure legends). The process of ILV formation observed in the absence of Vps4 and ATP is referred to as ‘passive’. A second fluorescent dextran (M_r_ ~ 10 kDa, AlexaFluor 488 labelled) can be added to this GUV population, followed by 20 minutes incubation with Vps4 and ATP.MgCl_2_. Newly formed ILVs in these conditions will contain blue and green fluorescence. The process of ILV formation driven by ATP hydrolysis by Vps4 is referred to as ‘active’. Both passive and active formation of ILVs, induced by the action of ESCRTs, are observed in GUVs (**Figure 2Bii**).

**Figure 2:**
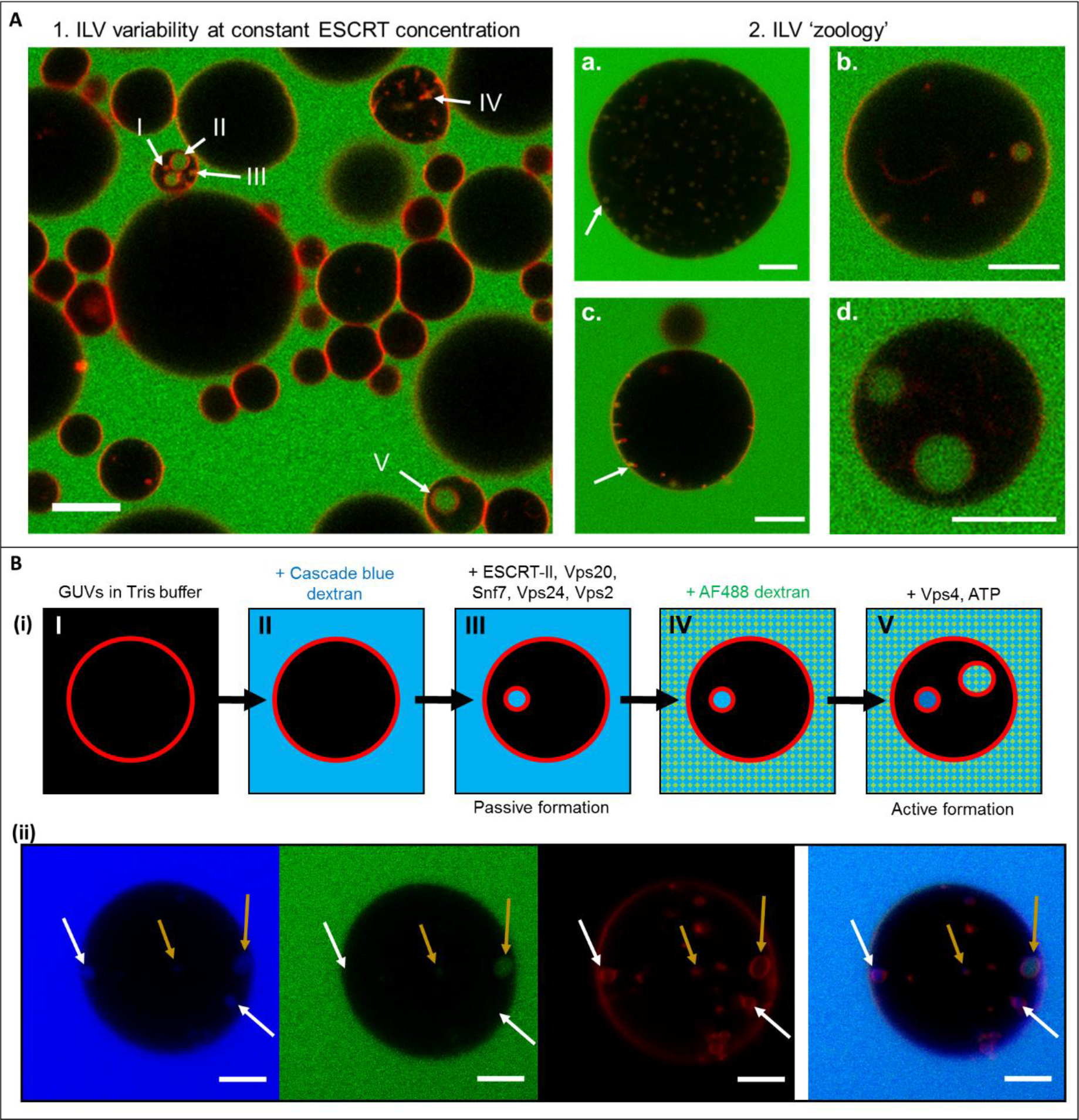
**A.1**: Confocal microscope image of a cross section (3.1 μm) through a number of GUVs incubated with ESCRT-II, Vps20, Vps24, Vps2 (10 nM) and Snf7 (50 nM). A number of different ILV sizes are observed in the field of view (I-V). Red: Rhodamine-PE, Green: AlexaFluor-488-labelled dextran. Scale bar 10 μm). **A.2**: ILV morphology, a range of commonly observed ILV states. **a.** Many similarly sized ILVs plus buds, **b.** Few ILVs of varying dimensions, **c.** ‘Stalled’ ILV buds with few or no co-incident free-floating ILVs, **d.** Unusually large ILVs ~5 μm diameter. Red: Rhodamine-PE, Green: AlexaFluor-488-labelled dextran. All scale bars 10 μm. **B(i):** Schematic of the ILV formation procedure. Cascade blue labelled dextran was added to GUVs in Tris buffer (**II**), immediately followed by ESCRT-II, Vps20, Vps24, Vps2 (10 nM) and Snf7 (50 nM) (**III**). After a 20 minute incubation period, AlexaFluor 488-labelled dextran was added, immediately followed by Vps4 (10 nM) and ATP.MgCl_2_ (1 μM). Thereby, ILVs containing only cascade blue dextran can be identified as having formed before the addition of Vps4 and ATP, and those that contain both dyes, must have formed after addition. **B(ii):** Confocal image of a GUV containing ILVs formed by active (ATP driven) and passive (no ATP) processes. Cascade Blue-labelled dextran (blue channel) was added immediately before incubation with ESCRT-II, Vps20, Vps24, Vps2 (10 nM) and Snf7 (50 nM). AlexaFluor488-labelled dextran (green channel) was added immediately before the addition of Vps4 (10 nM), and ATP.MgCl_2_ (1 μM) followed by a further period of incubation. White arrows indicate passively formed ILVs, yellow arrows indicate actively formed ILVs Membranes labelled with lissamine-rhodamine-PE (red channel). 40x objective, scale bar = 10 μm.

### ILV formation efficiency reaches a maximum for a specific molar ratio of Snf7

Snf7 oligomerisation is central to ESCRT-III activity, with more Snf7 units present in the active complex than other ESCRTs,^35^ but the overall stoichiometry, and its relationship to activity, is currently not well understood.

**Figure 3.**
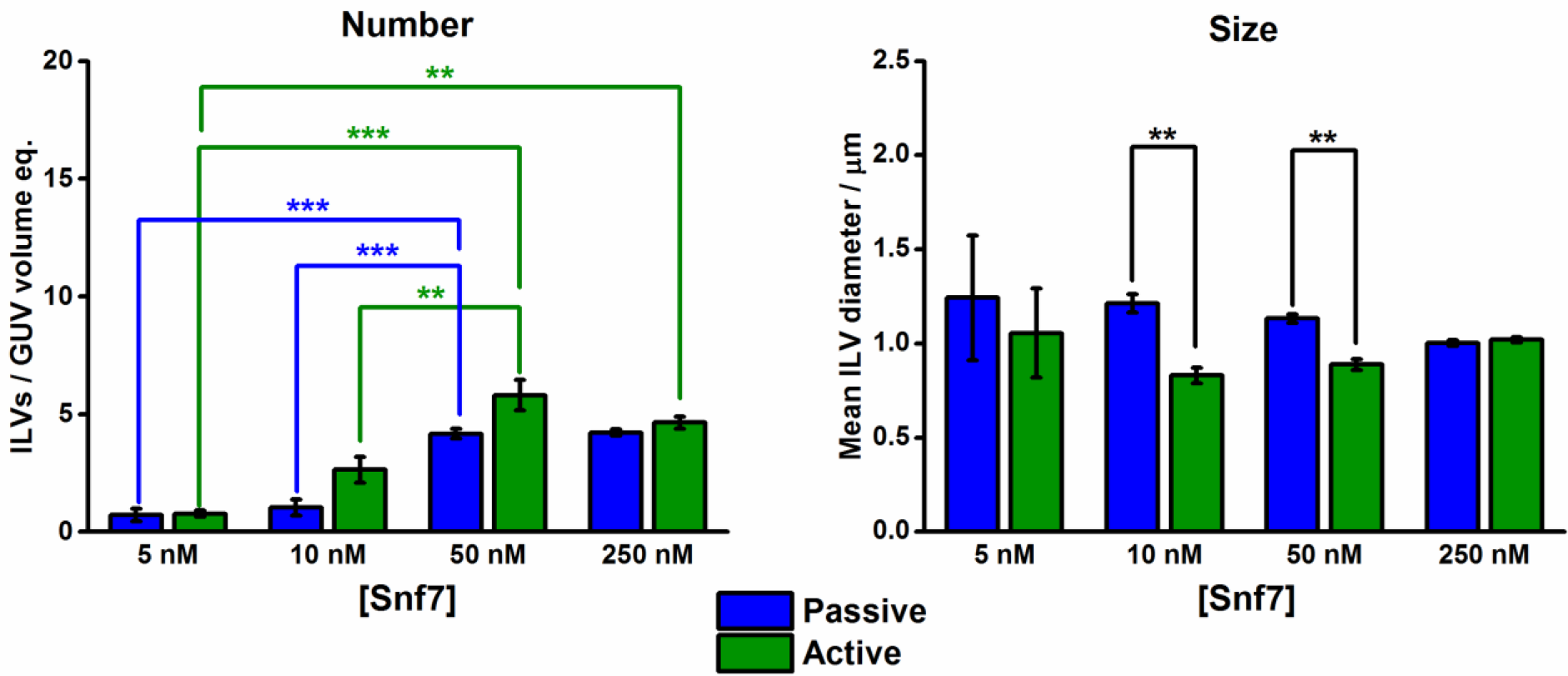
Snf7 stoichiometry bulk phase encapsulation assay. ILV counting and characterisation by confocal microscopy. ILVs containing fluorescent dye in their lumens after the addition of ESCRT proteins. **Number**: Number of ILVs per GUV volume equivalent; the number of ILVs observed per volume of a 20 μm diameter sphere. **Size**: Mean diameter of the observed ILVs. **Blue**: ‘Passive’ formation. ILVs containing cascade blue-dextran, after simultaneous addition of ESCRT-II, Vps20, Vps24, and Vps2 (10 nM), and Snf7 at the concentration stated on the x-axis, but before the addition of 10 nM Vps4 and 1 μM ATP.MgCl2. **Green**: ‘Active’ formation. ILVs containing AlexaFluor488-dextran observed after the addition of 10 nM Vps4 and 1 μM ATP.MgCl2. Osmotically relaxed vesicles were used in all experiments. Error bars are the standard error of the mean. Each data point is averaged from n=3 independent experiments, each containing 100+ independent GUVs. Significance testing: one-way ANOVA with Bonferroni test, one-tailed. No ESCRT control background rate = 0.067 ILVs per unit volume (standard error: 0.016).

Bulk-phase encapsulation assays are performed at a range of Snf7 concentrations (5-250 nM), and maintain a fixed concentration (10 nM) of each of the remaining ESCRT-II and ESCRT-III components (**Figure 3**). For both passive and active processes, 50 nM Snf7 provides the greatest activity in terms of number of ILVs observed, with no improvement at 250 nM. Interestingly, this is comparable to Snf7 being present at approximately up to five times the concentration of other ESCRT-III components in yeast.^29,35^

A similar number of ILVs form during the passive and active processes, which may indicate ‘recycling’ of the membrane bound ESCRT components once, and their reassembly into a similar number of active complexes. However other explanations are plausible and this may be complicated by changes and variability in GUV membrane tension in the second “active” round of ILV formation. ILV size is also found to be generally consistent across the full concentration range, with the only significant differences being found between active and passive processes at the 10 nM and 50 nM Snf7 concentrations, where ATP-fuelled “active” formation favours slightly smaller ILVs, on average. Below 10 nM and above 50 nM Snf7 may be too little or too much protein with respect to other ESCRT-III components to produce fully competent complexes.

### Membrane tension is the dominant factor in ILV formation efficiency and size. Protein stoichiometry effects are minimal

Vps20 bridges ESCRT-III to ESCRT-II and is involved in the nucleation of Snf7 oligomers onto endosomal membranes, *in vivo*.^27,36^ Vps20 possesses an N-terminal myristoylation that facilitates binding to membranes in general,^37^ but a transient association with the ESCRT-II subunit Vps25 is required to induce a conformational change that leads to the recruitment of Snf7.^27,37^ Nucleation is a critical step in the formation of ILVs, therefore Vps20 stoichiometry was examined for its influence on ILV size and number. Varying the Vps20 concentration relative to that of other ESCRT-III components should provide different numbers of potential nucleation sites for ESCRT-III assembly. A higher number of nucleation sites should result in a similarly higher number of ILVs, provided that there are sufficient other ESCRT-III components to form functional assemblies at each of these sites. The ‘dilution’ of the available ESCRT-III components across a larger number of nucleation sites may also result in smaller Snf7 assemblies, which may give rise to smaller ILVs, if there is a relationship between the size of the complex and the resulting ILV. Furthermore, these experiments are performed with both high and low membrane tension GUVs, where low tension GUVs are subjected to an osmotic relaxation protocol, to assess the influence of membrane mechanics.

**Figure 4.**
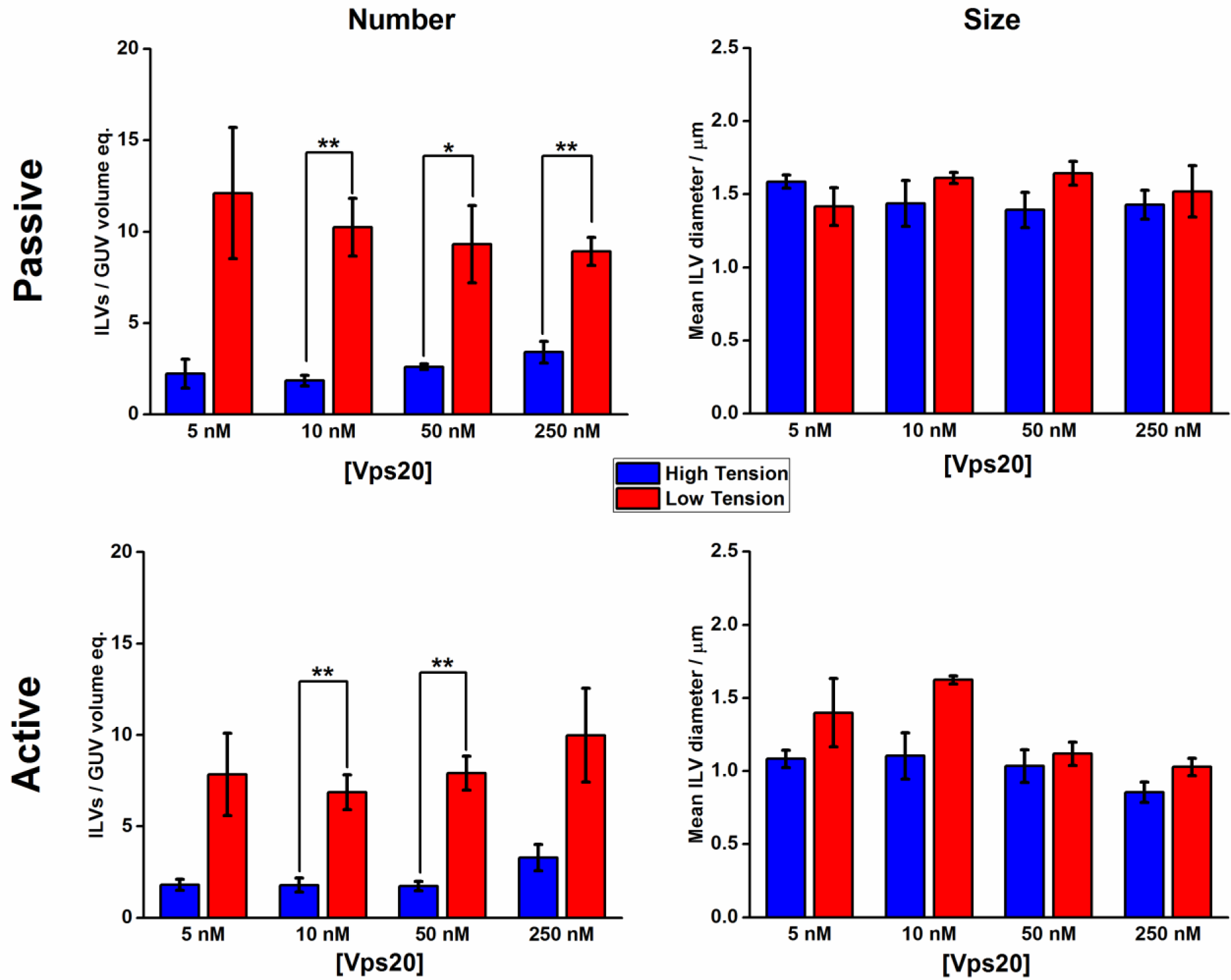
Vps20 stoichiometry bulk phase encapsulation assay. ILV counting and characterisation by confocal microscopy. ILVs containing fluorescent dye in their lumens after the addition of ESCRT proteins. **Number**: Number of ILVs per GUV volume equivalent. **Size**: Mean diameter of the observed ILVs. **Blue**: High membrane tension GUVs, **Red**: Low membrane tension GUVs. **Passive**: ILVs observed after simultaneous addition of ESCRT-II, Vps24, and Vps2 (10 nM), Snf7 (50 nM) and Vps20 at the concentration stated on x-axis, but before the addition of 10 nM Vps4 and 1 μM ATP.MgCl2. **Active**: After the addition of 10 nM Vps4 and 1 μM ATP.MgCl2. Error bars are the standard error of the mean. Each data point is averaged from n=3 independent experiments, each containing 100+ independent GUVs. Significance testing: one-way ANOVA with Bonferroni test, one-tailed. No ESCRT control background rate for low tension GUVs = 0.067 ILVs per unit volume (standard error: 0.016), for high tension GUVs = 0.054 ILVs per unit volume (standard error: 0.017).

Across a 50-fold variation in concentration, the relative stoichiometry of Vps20 to other ESCRT sub-units does not appear to have a significant effect on the number of ILVs observed (**Figure 4 ‘Number’**). While a reduction in membrane tension results in 3-4 times the number of ILVs formed per unit volume for both active and passive activity. Neither Vps20 stoichiometry nor membrane tension yield a significant effect on ILV size (**Figure 4 ‘Size’**).

In a recent model of ESCRT-III assembly,^21,28,38^ Vps24 and Vps2 bind to the growing end of the Snf7 filament providing a limiting signal to filament growth by terminating Snf7 spiral elongation. Vps2 recruits the ATPase Vps4 to the ESCRT-III assembly to recycle the proteins and allow for multiple rounds of ILV formation. However competing models have also recently suggested the formation of mixed Snf7, Vps2, Vps24 filaments where each protein has a different preferred curvature in the spiral assembly.^22^ In this experiment, we test how varying Vps24 stoichiometry influences the efficiency of ILV formation, which may occur either by premature termination of ESCRT-III filament growth or alternatively, by modulating the stored elastic energy in spiral ESCRT complexes.

**Figure 5.**
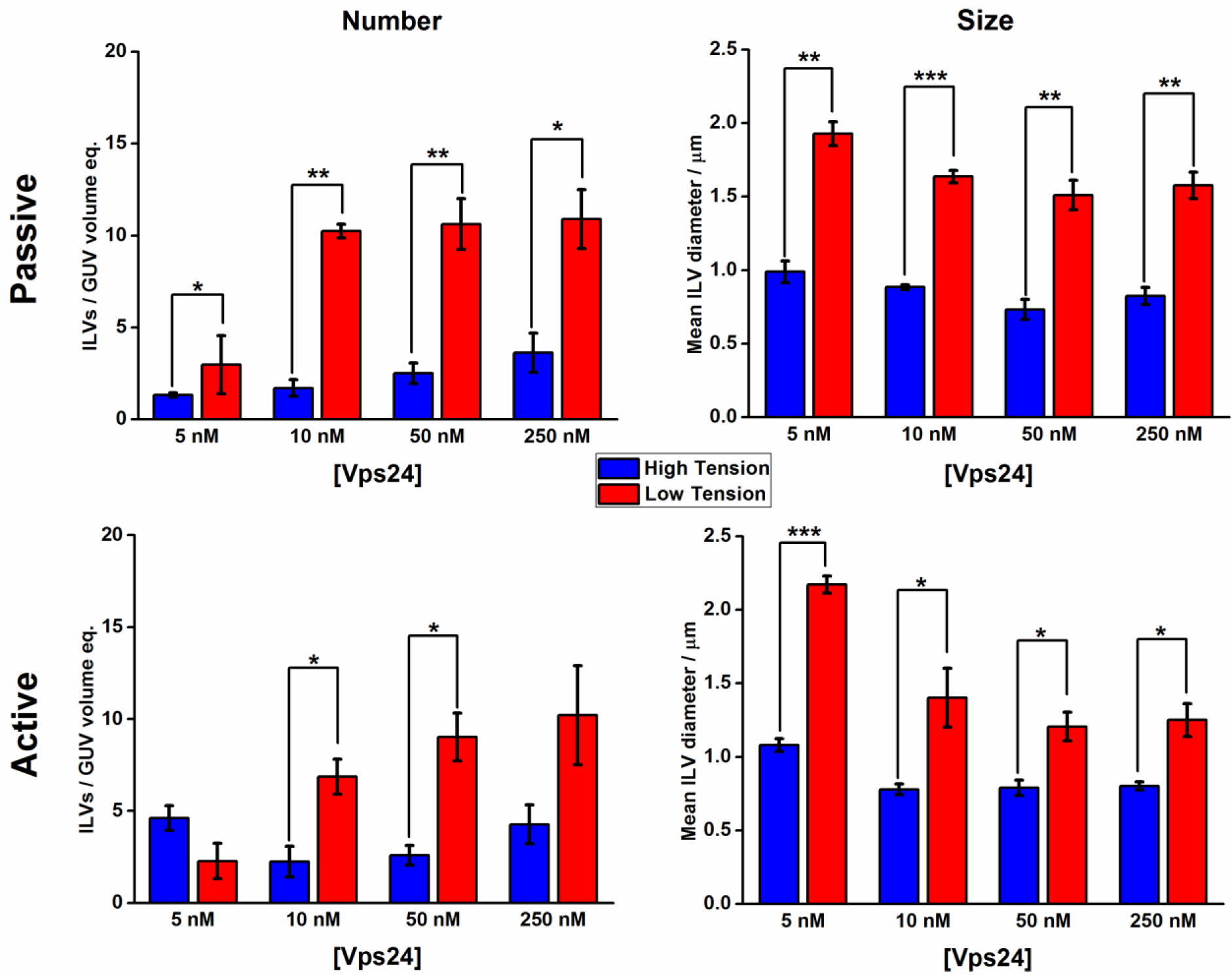
Vps24 stoichiometry bulk phase encapsulation assay. ILV counting and characterisation by confocal microscopy. ILVs containing fluorescent dye in their lumens after the addition of ESCRT proteins. **Number**: Number of ILVs per GUV volume equivalent. **Size**: Mean diameter of the observed ILVs. **Blue**: High membrane tension GUVs, **Red**: Low membrane tension GUVs. **Passive**: ILVs observed after simultaneous addition of ESCRT-II, Vps20, and Vps2 (10 nM), Snf7 (50 nM) and Vps24 at the concentration stated on x-axis, but before the addition of 10 nM Vps4 and 1 μM ATP.MgCl2. **Active**: After the addition of 10 nM Vps4 and 1 μM ATP.MgCl2. Error bars are the standard error of the mean. Each data point is averaged from n=3 independent experiments, each containing 100+ independent GUVs. Significance testing: one-way ANOVA with Bonferroni test, one-tailed. No ESCRT control background rate for low tension GUVs = 0.067 ILVs per unit volume (standard error: 0.016), for high tension GUVs = 0.054 ILVs per unit volume (standard error: 0.017).

Overall, we see a weak dependence of Vps24 stoichiometry on the efficiency of ILV formation in a similar fashion to what was observed for Vps20 (**Figure 5**), where the effects of membrane tension dominate the actions induced by stoichiometric changes of Vps24 with a 3-5 fold increase in ILV number in the low tension regime. However at the lowest Vps24 concentration (5 nM), there appears to be an inhibitory effect on ILV number in the low tension GUV population, but in high tension samples, no Vps24 concentration effect on number is observed. GUVs in the low tension regime also display an effect of Vps24 stoichiometry on the size of ILV produced, with the lowest Vps24 concentration producing significantly larger mean ILV diameter for both active and passive formation, suggestive that increased capping of the complex by Vps24 favours smaller ILV sizes. Furthermore, we see a significant increase in ILV size in the low tension regime in these experiments, indicative that lower mechanic resistance to membrane deformation facilitates more membrane, on average, being removed from the parent GUV in an individual ILV formation event.

Taken together, our data show that membrane tension plays a major role in regulating ESCRT-driven ILV formation.

### Membrane tension negatively regulates ILV formation by ESCRT-III

The deformation of a membrane during ESCRT-induced ILV budding would encounter greater mechanical resistance on higher tension membranes. The energetic input to achieve this may be derived from the assembly of ESCRT-III components, primarily Snf7 filaments, which are directly implicated in driving membrane budding. While the role of Vps4, a mechanoenzyme, has been suggested to be in the scission of ILV buds from their parent membrane through the disassembly of stabilising Snf7 filaments, some published results have suggested that ESCRT complexes only exert measurable forces on membranes during ATP-driven Vps4 activity.^26^

Our data reveals a strong, significant dependence of the efficiency of ILV formation on membrane tension, implying typical membrane tensions in GUVs are sufficient to compete with the energetic driving force of membrane deformation exerted by the ESCRT complex. Therefore, regardless of the mechanism or magnitude of the force exerted, there must exist a maximum membrane tension that is able to undergo ESCRT-induced ILV budding. A relationship between ILV size and membrane tension is also plausible as the magnitude of membrane deformation that can be achieved during budding may be modulated by tension. Any such effects resulting from resistance to membrane deformation will necessarily be concurrent with any effects resulting from transient membrane curvature. As assemblies of ESCRT-II and Vps20 are known to have an increased affinity for regions of membrane curvature,^27^ the transient curvature changes from membrane undulations that are more pronounced in low tension membranes may also enhance the affinity of ESCRT-II/ Vps20 complexes for the membrane.

Flicker spectroscopy experiments are used to quantify the distribution of membrane tensions (σ) in GUV populations in our low tension and high tension regimes, before and after incubation with ESCRT proteins (**Figure 4** and **Supp. Fig. S3.**).

**Figure 6:**
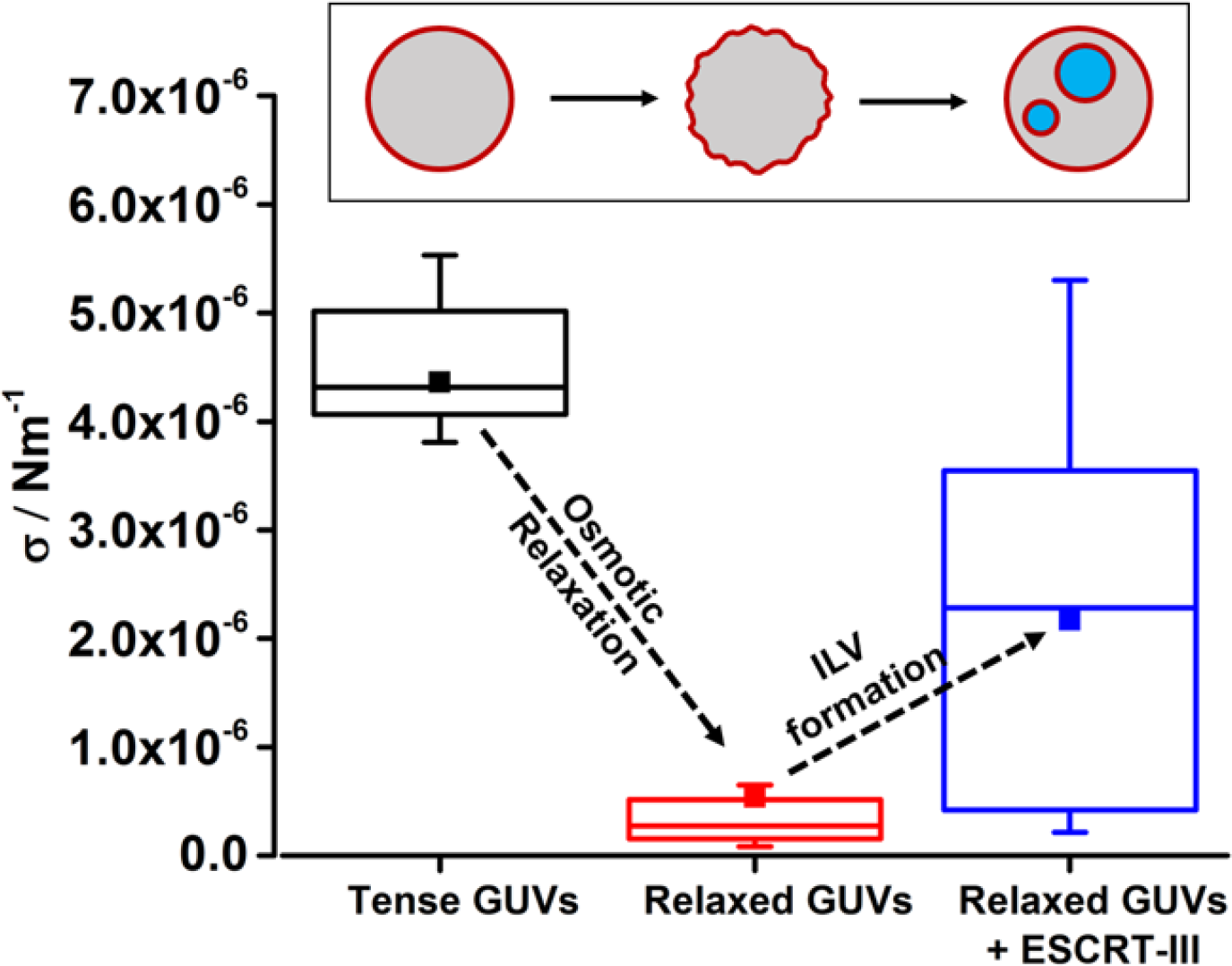
The distribution of membrane tension σ in GUV samples. ‘**Tense**’ GUVs were those that were not exposed to an osmotic gradient. The membrane tension of osmotically ‘relaxed’ GUVs was measured before (**Pre-protein**) and after (**Post-protein**) the addition of the ESCRT proteins to assess the effect of ILV formation on membrane tension. ■ – mean values. n=10.

GUVs are found to have high membrane tension following initial electroformation. Addition of the ESCRT complex resulted in a narrowing of the distribution of membrane tensions towards the high tension end of the initial distribution (**Supp. Fig. 3**). This implies that the lower tension vesicles within the initial population have increased their tension due to their interaction with ESCRTs. This is likely due to ILVs forming within the less tense vesicles of the high tension population and a significant number of GUVs where no ILVs have formed are observed in these samples, suggestive that their tension might be too high for the ESCRT complex to act upon them. Some GUVs in these samples are too tense to be quantified by the full flicker analysis as the data for higher *q* modes became noisy due to their small amplitude. Therefore the mean square amplitudes of the longest wavelength undulatory mode (n=3) is analysed as a proxy for membrane tension where smaller mean square amplitudes correspond to a higher membrane tension.^39^

The mechanical coupling between membrane tension and ESCRT function is most evident in the low tension regime. As expected, the osmotic relaxation procedure reduces the mean membrane tension by about an order of magnitude and also narrows the range of tension values across the population (**Figure 6 ‘Relaxed GUVs’**). Further flicker spectra are obtained after incubation with ESCRT proteins, using our standard procedure (50 nM Snf7, 10 nM ESCRT-II, Vps20, Vps24, Vps2, Vps4, 1 μM ATP.MgCl2, 20 minutes incubation), (**Figure 6. ‘Relaxed GUVs + ESCRT-III’**). The mean tension after ILV formation increases to an intermediate value between the ‘Tense’ GUVs and those that have been osmotically relaxed ‘Pre-protein’. This increase in membrane tension will act to make further ILV formation more difficult, providing a mechanism of negative feedback to self-regulate ESCRT activity. Notably, the ‘post-protein’ tension range does not reach the highest tension values observed in the initial electroformed GUV population, suggesting that in these cases, ILV formation is not limited by the membrane mechanics and further ILV formation is likely possible upon addition of more ESCRT-III or further active recycling of the existing components.

## Discussion

Here we have shown that ESCRT-III activity is regulated by the tension of the membrane substrate, using an *in vitro* model membrane system. Our work recapitulates ESCRT-mediated membrane budding and scission in the GUV system and shows that membrane tension significantly dominates over any attempt to control ILV formation by varying the ratio of the ESCRT-III assembly reaction. Increased membrane tension through removal of excess membrane surface area, provides a negative feedback mechanism that eventually suppresses membrane remodelling by ESCRT-III.

Based on a model where individual ESCRT constituents promote nucleation (Vps20), growth (Snf7) and termination (Vps24) of the ESCRT filaments, we hypothesized that variation of relative ratio between subunits may affect the efficiency of ILV formation and ILV size. Nonetheless, we observed minimal effects on ILV formation efficiency and size when varying Vps20 and Vps24 concentration across a 50-fold concentration range. This suggests that either: (i) membrane mechanics strongly dominates the effects of nucleation and termination in our experimental regime, or (ii) there is a narrow range of stoichiometric composition where functional ESCRT complexes form, giving rise to a weak dependence on the concentration of a single component. We consider the latter scenario to be most likely given that ILV formation in relaxed vesicles is not limited by membrane mechanics at the typical protein concentrations used (**Figure 6**).

Mechanistic models of ESCRT activity revolve around the formation of circular or spiral structures on membranes that drive membrane invagination,^20^ constricting and stabilising ILV bud necks^40^ prior to scission by disassembly of the oligomer.^41^ Nonetheless, membrane curvature drives the recruitment of specific ESCRT subunits^42^ and membrane undulations, amplified at low tension, have been implicated in promoting self-assembly of other curvature-generating proteins at the membrane.^43^ Hence, a more complete model for ESCRT-mediated remodelling activity includes the membrane as an active participant. Here, we have elucidated a regulatory role for membrane mechanics in preventing runaway over-activity of the ESCRT complex *via* a negative feedback mechanism.

ESCRTs perform a wide variety of membrane remodelling functions within the cell^12^ with some of these responding directly to changes in membrane tension, such as membrane abscission during cytokinesis, whereby relaxation of membrane tension promotes recruitment of ESCRT-III to the intercellular bridge.^44^ In the endocytic pathway, fusion of late endosomes would provide excess membrane area that lowers membrane tension and could therefore facilitate generation of internal vesicles by ESCRT-III, suggestive of a homeostatic relationship between endosomal fusion and formation of vesicles within the MVB. ESCRTs also play an important role in plasma and nuclear envelope membrane repair, where a loss in tension caused by membrane impairment could contribute to the rapid recruitment of the complex to the site of damage.

Membrane tension could provide a general means of regulating ESCRT activity. Plasma membrane tension has been reported in the low tens of μN m^-1^ range^45^, which is above the range of values observed in our GUV membranes. This may suggest that ESCRT remodelling activity is generally inhibited until local events at the membrane lower tension and promote ESCRT reactions. Cellular membranes lacking cytoskeleton support, such as blebs, have tensions typically in the μN m^-1^ range^45^, matching the higher end of our measured values. Accordingly, our data show a significant increase in efficiency of ESCRT activity (as measured by efficiency of ILV formation) associated with a change in mean tension from 4.8 μN m^-1^ to 0.6 μN m^-1^. These represent physiologically relevant values of membrane tensions supporting the inference of our model for ESCRT activity regulation by membrane mechanics not only *in vitro* but also *in vivo*.

Our primary interest is to develop the ESCRT proteins as a means of directing compartmentalisation in artificial cell systems, ultimately to enable dynamic control over their compartmentalised structures. While the fundamental ILV-forming activity of ESCRT-III in GUVs is well established, our intention is to determine means of exerting finer control over compartment parameters, such as size, number and cargo encapsulation. Membrane tension provides a promising control parameter to regulate ILV formation in GUVs and is suggestive of strategies that would allow multiple rounds of ILV formation to provide numerous distinct artificial organelles within GUV artificial cells.

## Methods

### Protein over-expression and purification

The plasmids for overexpression of Saccharomyces cervisiae ESCRT-III subunits Vps20, Snf7, Vps2 are a gift from James Hurley (addgene plasmids #21490, #21492, #21494). The yeast Vps24 gene is subcloned into a pRSET(A) expression vector, containing a His6-tag and Enterokinase protease cleavage site. All constructs are transformed into JM109 (DE3) (made in house). All proteins are expressed in autoinduction media, including trace elements (Formedium). 2 L of culture are grown in the presence of ampicillin (final concentration of 100 mg/L) for 3 hr at 37 °C and then the temperature is reduced to 25 °C overnight. The proteins (Vps20, Snf7, Vps2) are affinity purified using MBP resin (GE Healthcare) and eluted with 10 mM maltose. Vps24 is denatured in 5 M urea and applied to IMAC Sepharose resin charged with Ni2+ (GE Healthcare) and after washing is eluted over a 0 - 500 mM imidazole gradient. Elution fractions (Vps20, Snf7, Vps2) are cleaved using TEV protease (pET28-MBP-TEV is a gift from Zita Balklava & Thomas Wassmer; addgene plasmid # 69929),^30^ a second round of affinity chromatography to remove TEV and are finally purified by size exclusion chromatography (SEC; Superdex 75 column). Pooled fractions of ESCRT-III are maintained at concentrations not higher than 8 μM to avoid aggregation. The monomeric state of all proteins is verified using analytical SEC (Superdex 75 10/30 column; GE Healthcare) (**Figure S1**). Purified proteins are aliquoted in small volumes of around 50 μL for one-time use only to avoid damaging from freeze-thawing. The proteins were immediately flash frozen in liquid nitrogen and stored at −80° C until use. Full-length yeast Vps4 (a gift from James Hurley; addgene plasmid #21495) is purified from E. coli as a glutathione S-transferase fusion protein following the method in Wollert and Hurley 2009.^6^ The purified protein is concentrated to 20 μM, tested for ATPase activity and stored at −80 °C. Proteins are defrosted on ice prior to use.

### Electroformation of Giant Unilamellar Vesicles (GUVs)

Preparation of GUVs is performed using the electroformation method. A solution of lipids in chloroform (15 μL, 0.7 mM; (1-palmitoyl-2-oleoyl-*sn*-glycero-3-phosphocholine (POPC, 61.9 mol%), 1-palmitoyl-2-oleoyl-*sn*-glycero-3-phospho-L-serine (POPS, 10 mol%), cholesterol (25 mol%), 1,2-dioleoyl-*sn*-glycero-3-phospho-(1’-myo-inositol-3’-phosphate) (PI(3)P, 3 mol%), and lissamine-rhodamine-PE (0.1 mol%), all purchased from Avanti Polar Lipids Inc. Alabaster, Al. USA.) is applied as a thin layer over two Indium-tin oxide (ITO) glass slides (8-12 Ω/sq, Sigma-Aldrich) and dried under a stream of nitrogen gas. The slides are assembled into an electroformation chamber using a silicone rubber gasket to create a chamber between the opposing conductive surfaces, with strips of copper tape between the gasket and each conductive surface to serve as electrical contacts. The chamber is filled with sucrose solution (600 mM) and sealed with a silicone rubber plug. The chamber is placed in an oven at 60 °C and using a function generator, an AC voltage was applied to the system (3 V peak to peak, 10 Hz, sinusoidal waveform) for two hours, followed by gradual reduction in the frequency over approximately 10 minutes to facilitate the closure of nascent GUVs from the surface. The chamber is then allowed to cool to room temperature before the GUV suspension is harvested.

If desired, osmotic relaxation of membrane tension is achieved by overnight incubation at 4 °C of the resulting GUV suspension in Tris buffer (50 mM Tris, pH 7.4) adjusted to 10 mOsM higher than the sucrose electroformation solution (1:4, vol:vol Tris:GUV suspension) to facilitate the osmotic transport of water from the GUV lumens. Osmolarity is measured using an Advanced Instruments 3320 osmometer and adjusted by addition of NaCl.

### Confocal microscopy - ILV counting

Confocal microscopy is performed on a Zeiss LSM 880 inverted system, using a Plan-Apochromat 40x/1.4 Oil DIC M27 objective lens, NA = 1.4. 8-well glass bottomed imaging chambers (ibidi GmbH) are prepared by passivation of the interior glass surface by incubation overnight in 5 % Bovine serum albumin (BSA) solution, followed by copious rinsing with MilliQ water. A GUV suspension is diluted 1:4 with osmotically balanced Tris buffer, unless previously diluted with 10 % hyperosmotic Tris buffer as described previously for membrane relaxation. To ensure efficient mixing for each experiment, solutions are combined in an Eppendorf tube, followed by gentle agitation of the tube and transfer of the solution to the imaging chambers. Proteins are thawed over ice, before being added to the tubes such as to give final concentration of 10 nM ESCRT-II, Vps20, Vps24 and Vps2, and 50 nM Snf7, unless otherwise stated. Aliquots of proteins are used only once and never re-frozen and thawed again. Fluorescently labelled dextran (M_r_ ~ 5,000 Da, Cascade blue labelled) is then added, followed by 200 μL of GUV suspension. Imaging is then performed after 20 minutes of incubation. Once imaging is complete, a second fluorescent dextran (M_r_ ~ 10,000 Da, AlexaFluor 488 labelled) is added to each sample, followed by 10 nM Vps4 and 1 μM ATP.MgCl_2_ and further imaging is performed after a 20 minute incubation period. For the ILV counting experiments, a tile scanning technique with a 3.1 μm section thickness is employed to capture a cross sectional volume through a large number of GUVs. Only ILVs visibly containing encapsulated dextran dye are counted, indicating that they have formed after the addition of the proteins.

### Flicker spectroscopy

GUV membrane tension is quantified by flicker spectroscopy.^31–33^ This technique analyses the power spectrum of membrane undulations (mean square undulation amplitude 〈|*u*(*q*)|^2^〉 versus wavenumber *q* from image analysis of a confocal microscopy time series taken at the midpoint (equator) of the GUV. 1000 frames are taken per GUV at 1024×1024 pixel resolution, frame rate and pixel size varied on a case by case basis as the maximum magnification that could accommodate the whole GUV is used to maximize scan speed and these factors are accounted for in the MATLAB analysis software. Fitting 〈|*u*(*q*)|^2^〉 with the equation:

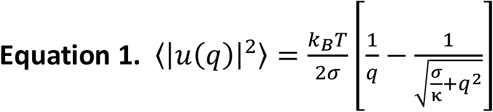

allows us to quantify the bending rigidity (κ) and tension (σ) of individual GUV membranes.^34^ The MATLAB program for GUV flicker analysis was kindly provided to us by Prof. Pietro Cicuta and co-workers at the University of Cambridge, UK. An example flicker spectrum is shown in supplementary **Figure S4**.

## Acknowledgements

This research is supported by the UK Engineering and Physical Sciences Research Council (EPSRC): EP/M027929/1 (PAB); EP/M027821/1 (BC). We would like to thank Dr Daniel Mitchell for the expression and purification of the Vps4 subunit. We would also like to thank Prof. Pietro Cicuta and Dr. Lucia Parolini of the University of Cambridge for sharing their MatLab programmes for flicker spectroscopy analysis and their assistance in its initial use.

